# Genome assembly of the rare and endangered Grantham’s camellia, *Camellia granthamiana*

**DOI:** 10.1101/2024.01.15.575486

**Authors:** Hong Kong Biodiversity Genomics Consortium, Jerome H.L. Hui, Ting Fung Chan, Leo L. Chan, Siu Gin Cheung, Chi Chiu Cheang, James K.H. Fang, Juan Diego Gaitan-Espitia, Stanley C.K. Lau, Yik Hei Sung, Chris K.C. Wong, Kevin Y.L. Yip, Yingying Wei, Sean T.S. Law, Wai Lok So, Wenyan Nong, Sean T.S. Law, Wenyan Nong, David T.W. Lau, Ho Yin Yip

**Affiliations:** School of Life Sciences, Simon F.S. Li Marine Science Laboratory, State Key Laboratory of Agrobiotechnology, Institute of Environment, Energy and Sustainability, The Chinese University of Hong Kong, Hong Kong, China; School of Life Sciences, State Key Laboratory of Agrobiotechnology, The Chinese University of Hong Kong, Hong Kong SAR, China; State Key Laboratory of Marine Pollution and Department of Biomedical Sciences, City University of Hong Kong, Hong Kong SAR, China; State Key Laboratory of Marine Pollution and Department of Chemistry, City University of Hong Kong, Hong Kong SAR, China; Department of Science and Environmental Studies, The Education University of Hong Kong, Hong Kong SAR, China; EcoEdu PEI, Charlottetown, PE, C1A 4B7, Canada; Department of Food Science and Nutrition, Research Institute for Future Food, and State Key Laboratory of Marine Pollution, The Hong Kong Polytechnic University, Hong Kong SAR, China; The Swire Institute of Marine Science and School of Biological Sciences, The University of Hong Kong, Hong Kong SAR, China; Department of Ocean Science, The Hong Kong University of Science and Technology, Hong Kong SAR, China; Science Unit, Lingnan University, Hong Kong SAR, China; School of Allied Health Sciences, University of Suffolk, Ipswich, IP4 1QJ, UK; Croucher Institute for Environmental Sciences, and Department of Biology, Hong Kong Baptist University, Hong Kong SAR, China; Department of Computer Science and Engineering, The Chinese University of Hong Kong, Hong Kong SAR, China; Sanford Burnham Prebys Medical Discovery Institute, La Jolla, CA, USA; Department of Statistics, The Chinese University of Hong Kong, Hong Kong SAR, China; Shiu-Ying Hu Herbarium, School of Life Sciences, The Chinese University of Hong Kong, Hong Kong SAR, China

## Abstract

The Grantham’s camellia (*Camellia granthamiana* Sealy) is a rare and endangered tea species that is endemic to southern China, and was first discovered in Hong Kong in 1955. Despite its high conservation value, genomic resources of *C. granthamiana* remain limited. Here, we present a chromosome-scale draft genome of the tetraploid *C. granthamiana* (2n = 4x = 60) using a combination of PacBio long read sequencing and Omni-C data. The assembled genome size is ∼2.4 Gb with most sequences anchored to 15 pseudochromosomes that resemble a monoploid genome. The genome is of high contiguity, with a scaffold N50 of 139.7 Mb, and high completeness with a 97.8% BUSCO score. Gene model prediction resulted in a total 76,992 protein-coding genes with a BUSCO score of 85.9%. 1.65 Gb of repeat content was annotated, which accounts for 68.48% of the genome. The Grantham’s camellia genome assembly provides a valuable resource for future investigations on its biology, ecology, phylogenomic relationships with other *Camellia* species, as well as set up a foundation for further conservation measures.

## Introduction

*Camellia* is a large genus in the family Theaceae with more than 230 described species (POWO, 2021). While some camellias are well-known for their ornamental and economical values as tea and woody-oil producing plants that derived into tens of thousands of cultivars (Wang et al., 2021), more than 60 *Camellia* species were regarded as globally threatened due to natural habitat fragmentation or loss and small population size (Beech et al., 2017). The Grantham’s camellia (*Camellia granthamiana*) (Figure 1A) is a rare species once discovered in Hong Kong and named after Sir Alexander Grantham and is narrowly distributed in Hong Kong and Guangdong, China (Beech et al., 2017). It is listed as vulnerable in the IUCN Red List and recorded as endangered in the China Plant Red Data Book (Fu & Chin, 1992). In Hong Kong, the Grantham’s camellia is a protected species by law and has been actively being propagated and reintroduced to the wild by the Agriculture, Fisheries and Conservation Department (Hu, 2003).

**Figure 1.**
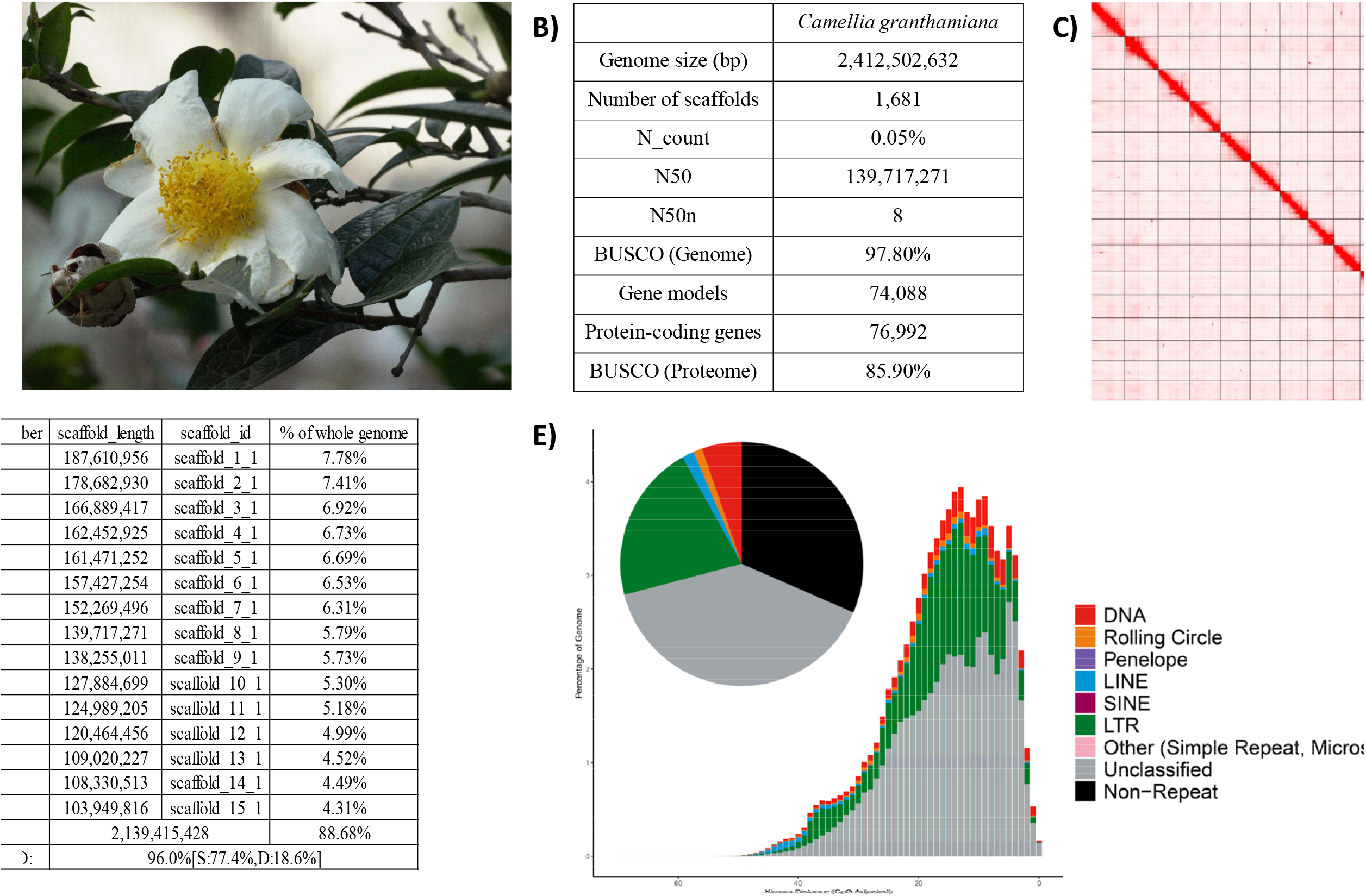
Genomic information of *Camellia granthamiana*. **A)** Picture of *Camellia granthamiana*; **B)** Summary of genome statistics; **C)** Omni-C contact map of the genome assembly; **D)** Information of 15 pseudochromosomes; **E)** Pie chart (Top) and repeat landscape plot (bottom) of repetitive elements in the genome.

## Context

In view of the high conservation value of Grantham’s camellia, several molecular studies have been previously conducted. They include sequencing the chloroplast genomes of *C. granthamiana* (Jiang et al., 2019; Li et al., 2018), using pan-transcriptomes to reconstruct the phylogeny of over a hundred of *Camellia* species (Wu et al., 2022), and population genetics study (Chen et al., 2023). However, nuclear genomic resources of *C. granthamiana* remain lacked. While most *Camellia* species possess a karyotype of 2n = 30, *C. granthamiana* is one of the exceptions with a karyotype of 2n = 4x = 60 (Huang et al., 2013; Kondo et al., 1977).

In Hong Kong, *C. granthamiana* was chosen as one of the listed species for sequencing in the Hong Kong Biodiversity Genomics Consortium (a.k.a. EarthBioGenome Project Hong Kong), which is formed by investigators from eight publicly funded universities. Herein, we report the genome assembly of *C. granthamiana* which can serve as a solid foundation for further investigations of this rare and endangered species.

## Methods

### Sample collection and high molecular weight DNA extraction

Fresh leaf tissues were sampled in transplanted individual on the campus of the Chinese University of Hong Kong. High molecular weight (HMW) genomic DNA was isolated from 1g leaf tissues using a CTAB pretreatment followed by NucleoBond HMW DNA kit (Macherey Nagel Item No. 740160.20). Briefly, the tissues were ground with liquid nitrogen and digested in 5 mL CTAB buffer (Doyle & Doyle, 1987) with an addition of 1% polyvinylpyrrolidone (PVP) for 1 h. The lysate was treated with RNAse A, followed by an addition of 1.6 mL of 3M potassium acetate and two round of chloroform:IAA (24:1) washes. The supernatant was transferred to a new 50 mL tube using a wide-bore tip. H1 buffer from the NucleoBond HMW DNA kit was added to the supernatant for a total volume of 6 mL mixture, from which the DNA was isolated by following the manufacturer’s protocol. After the DNA was eluted with 60 μL elution buffer (PacBio Ref. No. 101-633-500), quality check was carried out with NanoDrop^™^ One/OneC Microvolume UV-Vis Spectrophotometer, Qubit^®^ Fluorometer, and overnight pulse-field gel electrophoresis.

### Pacbio library preparation and sequencing

The qualified DNA was sheared with a g-tube (Covaris Part No. 520079) for 6 passes of centrifugation at 1,990 x *g* for 2 min and was subsequently purified with SMRTbell^®^ cleanup beads (PacBio Ref. No. 102158-300). 2 μL sheared DNA was taken for fragment size examination through overnight pulse-field gel electrophoresis. Two SMRTbell libraries were constructed with the SMRTbell® prep kit 3.0 (PacBio Ref. No. 102-141-700), following the manufacturer’s protocol. The final library was prepared with the Sequel^®^ II binding kit 3.2 (PacBio Ref. No. 102-194-100) and was loaded with the diffusion loading mode with the on-plate concentration set at 90 pM on the Pacific Biosciences SEQUEL IIe System, running for 30-hour movies to output HiFi reads. In total, three SMRT cells were used for the sequencing. Details of the resulting sequencing data are summarized in Supplementary Information 1.

### Omni-C library preparation and sequencing

Nuclei was isolated from 3 g fresh leaf tissues ground with liquid nitrogen using the PacBio protocol modified from Workman et al. (2018) (https://www.pacb.com/wp-content/uploads/Procedure-checklist-Isolating-nuclei-from-plant-t issue-using-TissueRuptor-disruption.pdf). The nuclei pellet was snap-frozen with liquid nitrogen and stored at -80 °C. Upon Omni-C library construction, the nuclei pellet was resuspended in 4 mL 1X PBS buffer and processed with the Dovetail® Omni-C® Library Preparation Kit (Dovetail Cat. No. 21005) by following the manufacturer’s procedures. The concentration and fragment size of the resulting library was assessed by Qubit^®^ Fluorometer and TapeStation D5000 HS ScreenTape, respectively. The qualified library was sent to Novogene and sequenced on an Illumina HiSeq-PE150 platform. Details of the resulting sequencing data are summarized in Supplementary Information 1.

### Genome assembly and gene model prediction

*De novo* genome assembly was first proceeded with Hifiasm (Cheng et al., 2021) and then was processed with searching against the NT database with BLAST to remove possible contaminations using BlobTools (v1.1.1) (Laetsch & Blaxter, 2017). Subsequently, haplotypic duplications were removed according to the depth of HiFi reads using “purge_dups” (Guan et al., 2020). Proximity ligation data from Omni-C were used to scaffold the assembly with YaHS (Zhou et al., 2022).

Gene models were trained, predicted and updated by funannotate (Palmer & Stajich, 2020) with the following parameters “--repeats2evm --protein_evidence uniprot_sprot.fasta --genemark_mode ET --optimize_augustus --organism other --max_intronlen 350000”. Seven RNA sequencing data were downloaded from NCBI (SRA Accessions: SRR16685015, SRR16685016, SRR16685017, SRR19086193, SRR19266768, SRR24821546, and SRR24821547) and aligned to the repeat soft-masked genome using Hisat2 to run the genome-guided Trinity (Grabherr et al., 2011), from which 289,554 transcripts were derived. The Trinity transcript alignments were converted to GFF3 format and used as input to run the PASA alignment to generate PASA models trained by TransDecoder, which were screened using Kallisto TPM data. The PASA gene models were used to train Augustus in the funannotate-predict step. The predicted gene models were combined from various prediction sources, including GeneMark, high-quality Augustus predictions (HiQ), pasa, Augustus, GlimmerHM and snap, and were integrated to produce the annotation files with Evidence Modeler. UTRs were further captured in the funannotate-update step using PASA.

### Repeat annotation

The annotation of transposable elements (TEs) were performed by the Earl Grey TE annotation pipeline (version 1.2, https://github.com/TobyBaril/EarlGrey) (Baril et al., 2022).

### Macrosynteny analysis

The longest gene transcripts from the predicted gene models of *Camellia granthamiana* and *Camellia sinensis* (accession number: GWHASIV00000000; Zhang et al., 2021) were used to retrieve orthologous gene pair with reciprocal BLASTp (e-value 1e-5) using diamond (v2.0.13) (Buchfink et al., 2021). The BLAST output was passed to MCScanX (Wang et al., 2012) to infer macrosynteny of the pseudochromosomes between *C. granthamiana* and *C. sinensis* with default parameters.

## Results and discussion

### Genome assembly of Camellia granthamiana

A total of 54.4 Gb HiFi reads was yielded from PacBio sequencing with an average length of 10,731 bp (Table 1; Supplementary Information 1). Together with 233.8 Gb Omni-C data, the genome of *Camellia granthamiana* was assembled with a final size of 2,412.5 Mb, from which 79.87% of the sequences were anchored into 15 pseudochromosomes (Figure 1B-1D; Supplementary Information 1). The scaffold N50 was 139.7 Mb and the BUSCO score was 97.8% (Figure 1B; Table 1). Gene model prediction yielded a total of 76,992 protein-coding genes with a mean length of 301 bp and BUSCO score of 85.9%.

Repeat content analysis annotated 1.65 Gb of transposable elements (TEs), comprising 68.48% of the *C. granthamiana* genome. Among the classified TEs, LTR retransposons accounted for the largest proportion (20.99%), followed by DNA transposons (5.30%), LINE (1.60%) and Rolling-circle transposons (1.21%) (Figure 1D; Table 2). The large proportion of repeat content in the *C. granthamiana* genome is comparable to other tea species, such as the Tieguanyin cultivar of *Camellia sinensis* (78.2%) (Zhang et al., 2021), wild oil-Camellia *Camellia oleifera* (76.1%) (Lin et al., 2022), and *Camellia chekiangoleosa* (79.09%) (Shen et al., 2022).

### Macrosynteny between Camellia granthamiana and Camellia sinensis

Macrosynteny analysis revealed a 1-to-1 pair relationship between the 15 pseudochromsomes of *C. granthamiana* and that of *C. sinensis* (Figure 2). This indicates that the assembled 15 pseudochromosomes resemble a monoploid genome of the tetraploid *C. granthamiana*.

**Figure 2.**
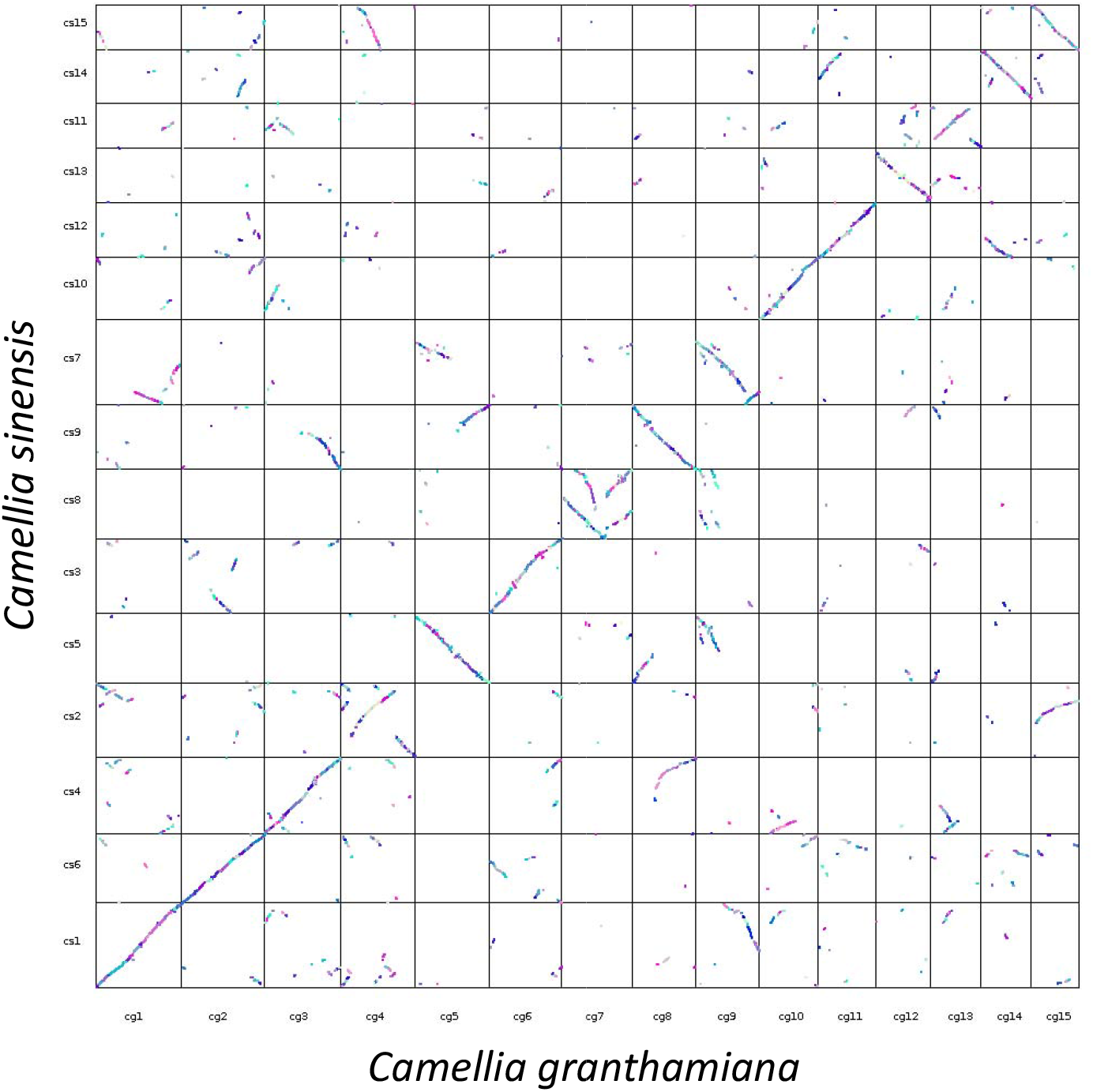
Macrosynteny dot plot between *Camellia granthamiana* and *Camellia sinensis*.

## Conclusion and future perspective

This study presents the first *de novo* genome assembly of the rare and endangered *C. granthamiana*. This valuable genome resource is of great potential for the use in future studies on the conservation biology of the Grantham’s camellia, its relationship with other *Camellia* species from a phylogenomic perspective and further investigations on the biosynthesis of secondary metabolites of tea species.

## Data validation and quality control

For HMW DNA and Pacbio library samples, NanoDrop^™^ One/OneC Microvolume UV-Vis Spectrophotometer, Qubit^®^ Fluorometer, and overnight pulse-field gel electrophoresis were used for quality control. The quality of Omni-C library was checked by Qubit^®^ Fluorometer and TapeStation D5000 HS ScreenTape.

During genome assembly, BlobTools (v1.1.1) (Laetsch & Blaxter, 2017) was employed to remove possible contaminations (Supplementary Information 2). The resulting genome assembly was run with Benchmarking Universal Single-Copy Orthologs (BUSCO, v5.5.0) (Manni et al., 2021) with the Viridiplantae dataset (Viridiplantae Odb10) to assess the completeness of the genome assembly and gene annotation.

## Disclaimer

The genomic data generated in this study was not fully haploptype-resolved for a tetraploid genome and the genome heterozygosity was not assessed.

## Data availability

The final genome assembly in this study was submitted to NCBI under accession number JAXFYN000000000. The raw reads generated were deposited in the NCBI database under under the SRA accessions SRR26895683 and SRR26909376. The genome annotation files were uploaded to Figshare (https://figshare.com/s/4b13376ad27ae0647fd1).

## Authors’ contribution

JHLH, TFC, LLC, SGC, CCC, JKHF, JDG, SCKL, YHS, CKCW, KYLY and YW conceived and supervised the study. DTWL collected the sample materials; STSL and WLS performed DNA extraction, library preparation and genome sequencing; HYY facilitated the logistics of samples; WN performed genome assembly, gene model prediction and genome quality check analyses; STSL carried out macrosynteny analysis.

---

**Table 1.** Genome statistics and sequencing information.

**Table 2.** Summary of classified transposable elements in the genome.

**Supplementary Information 1**. Summary of genomic sequencing data.

**Supplementary Information 2**. Genome assembly QC and contaminant/cobiont detection.

