## Supplementary material for "Genome assembly of the rare and endangered Grantham’s camellia, *Camellia granthamiana*": EBPHK_Camellia Supplementary Information.v2.docx

**Supplementary Information 1.** Genome sequencing information.

| **Liabrary** | **Reads** | **Bases** | **accession number** |
| --- | --- | --- | --- |
| PacBio HiFi | 5,071,365 | 54,421,045,547 | SRR26895683 |
| Omnic | 1,558,845,532 | 233,826,829,800 | SRR26909376 |

**Supplementary Information 2.** Genome assembly QC and contaminant/cobiont detection

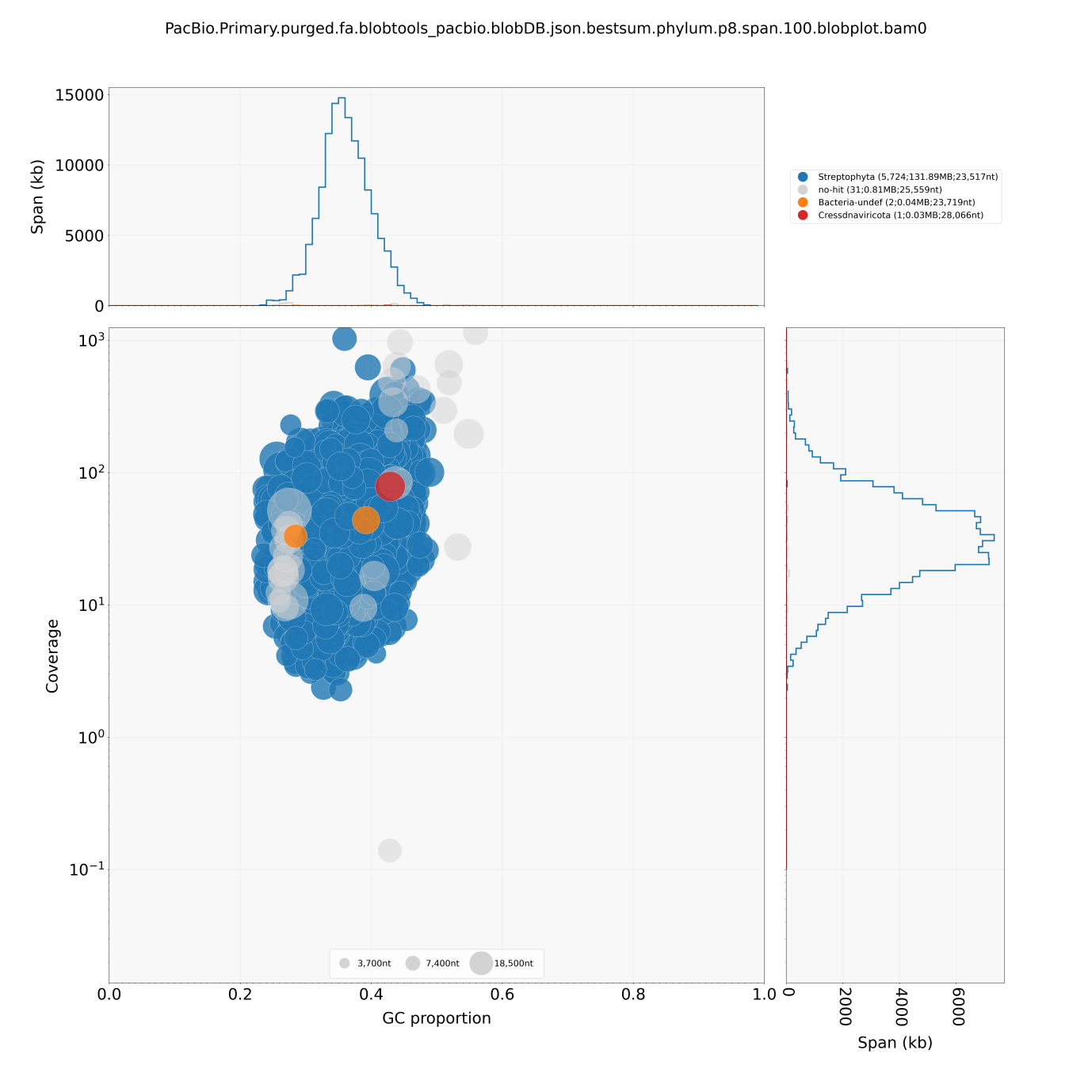

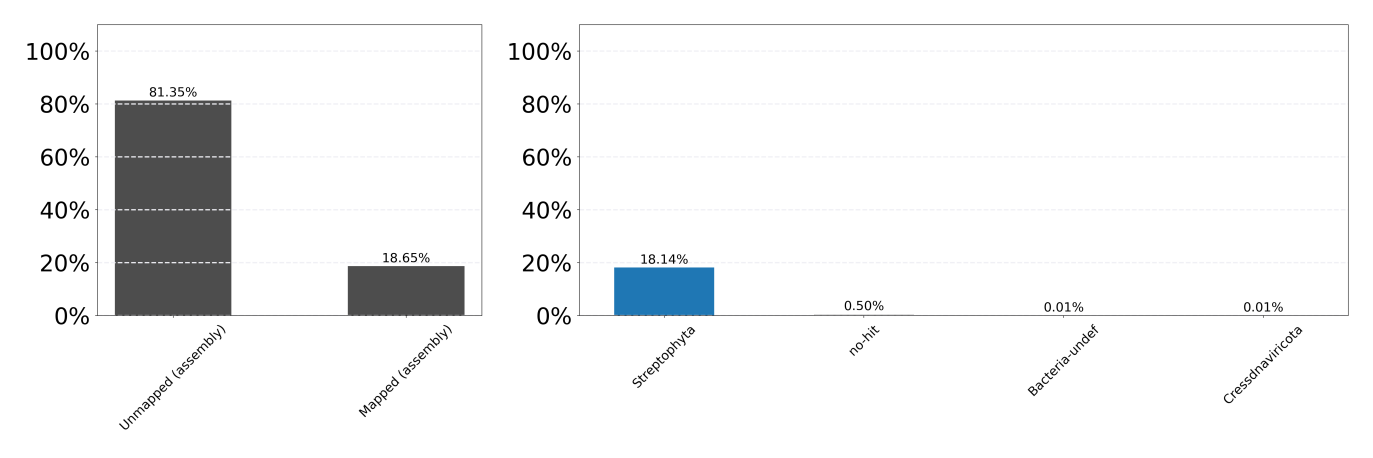
